# Hierarchical small molecule inhibition of MYST acetyltransferases

**DOI:** 10.1101/2025.04.25.650653

**Authors:** Xuemin Chen, Alexandra Castroverde, Minervo Perez, Ronald Holewinski, Kiall F. Suazo, Rashmi Karki, Thorkell Andressen, Benjamin A. Garcia, Jordan L. Meier

**Affiliations:** Chemical Biology Laboratory, National Cancer Institute, Frederick, Maryland, 21702, United States; Protein Characterization Laboratory, Frederick National Laboratory for Cancer Research, Leidos Biomedical Research, Frederick, Maryland, 21701, United States; Department of Biochemistry and Molecular Biophysics, Washington University School of Medicine, St. Louis, Missouri, 63110, United States

## Abstract

MYST lysine acetyltransferases (KATs) is a class of epigenetic enzymes critical for cellular function that constitute an emerging therapeutic target in cancer. Recently, several drug-like MYST inhibitors have been reported that show promise in a variety of preclinical models as well as in clinical trials of breast cancer. However, the comparative properties of these small molecules remains to be directly assessed. Here we apply an integrated profiling strategy to systematically define the potency and selectivity of drug-like MYST KAT inhibitors. First, we use optimized chemoproteomic profiling and histone acetylation biormarkers to study the industry-developed KAT inhibitor PF-9363. This reveals dose-dependent engagement of native KAT complexes, with hierarchical inhibition following the order KAT6A/B > KAT7 >> KAT8 > KAT5. Next, we demonstrate how PF-9363’s ability to disrupt capture of MYST complex members in chemoproteomic experiments can be leveraged to identify new candidate members of these complexes, including the transcription factor FOXK2. Applying insights from these studies to WM-8014, WM-1119 and WM-3835, which have been extensively applied in the literature as MYST probes, highlights unexpected cross-inhibition and suggests a new framework for how these small molecules and biomarkers may be applied to differentiate KAT6A/B and KAT7-dependent phenotypes. Finally, we benchmark the activity of PF-9363 in the NCI-60 cell line screen, providing evidence that it can inhibit the growth of cell lines that are resistant to other epigenetic inhibitors by engaging the essential MYST enzyme KAT8 at high concentrations. Collectively, our studies indicate the potential for MYST KAT inhibitors to exhibit dose-dependent target engagement reminiscent of kinase inhibitors and specify assays and biomarkers for facile monitoring of selective and hierarchical effects.

## Introduction

Lysine acetylation is a prevalent post-translational modification (PTM) catalyzed by lysine acetyltransferase (KAT) enzymes.^1^ The MYST family of KATs includes five members: KAT5, KAT6A, KAT6B, KAT7, and KAT8 (**Fig. 1A**).^2^ These genes are conserved from yeast to humans and are required for many critical nuclear functions, including transcription and DNA repair. The human disease relevance of these enzymes is exemplified by KAT6A and KAT6B, two closely related MYST paralogues whose translocation can form gene fusions capable of driving acute myeloid leukemia (AML).^3, 4^ Furthermore, KAT6A amplifications occur in 12-15% of breast cancers, and are hypothesized to facilitate tumor growth by regulating estrogen receptor (ER) signaling. ^5, 6^ These associations have motivated the development of selective small molecule inhibitors of MYST enzymes. Early efforts yielded WM-1119 (**Fig. 1B**), a drug-like KAT6A/B inhibitor capable of inducing cell cycle arrest and senescence in MYC-driven lymphoma models.^7, 8^ More recently, an optimized arylsulfonamide benzisoxazole PF-9363 (**Fig. 1B**) has been shown to disrupt ER-mediated gene expression and demonstrate in vivo efficacy in KAT6A-dependent solid tumor xenografts.^9^ An analogue of PF-9363 being tested in clinical trials has demonstrated promising pharmacokinetics, modulation of histone acetylation, and antitumor activity in ER+ breast cancer patients.^10^

**Figure 1.**
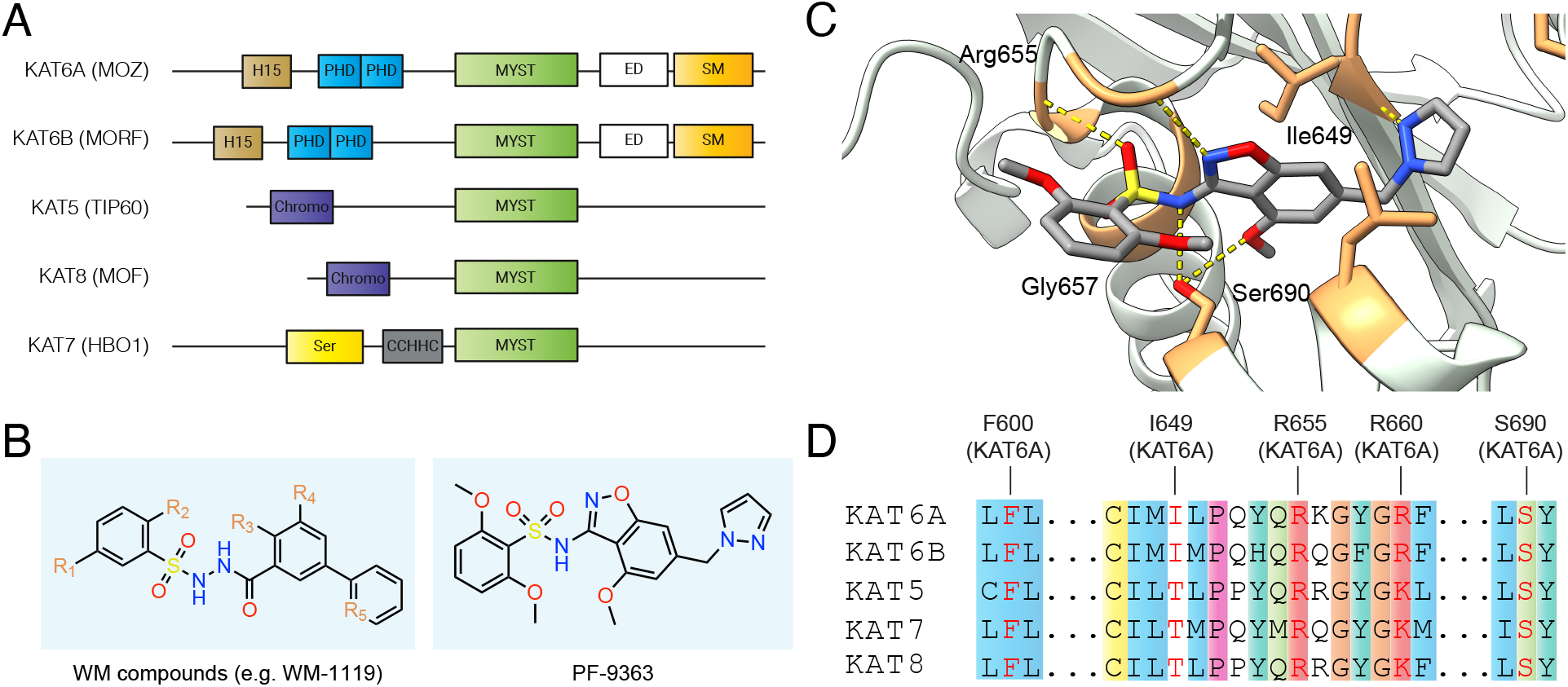
(A) Domain architecture of MYST lysine acetyltransferase family. H15: H15 domain; PHD: plant homeodomain-linked zinc finger; MYST: MYST type domain with KAT activity; ED: glutamate/aspartate-rich region; SM: serine/methionine-rich domain; Chromo: chromodomain; Ser: serine-rich domain; CCHHC: Zinc finger CCHHC-type. (B) Chemical structures of drug-like MYST inhibitors of the sulfonohydrazide (WM compounds) and benzisoxazole sulfonamide (PF-9363) class. (C) Crystal structure of PF-9363 bound to KAT6A catalytic domain. PDB code: 8DD5. (D) Sequence alignment of ligand-binding residues in MYST enzymes. Active site residues interacting with PF-9363 are highlighted in red. Residues were shaded based on the default color scheme for ClustalX if >60% matched the following characteristics: hydrophobic (blue); positive (red); polar (green); cysteine (yellow); proline (pink); aromatic (aqua); glycine (orange).

Nature directs MYST catalytic activity to specific histone residues using protein-protein interactions.^11, 12^ In contrast, selective small molecule inhibitors must differentiate the nearly identical active sites of this family (**Fig. 1C-D**). Structural studies indicate a highly conserved acetyl-CoA binding mode, with the cofactor engaging similar active site residues in each enzyme.^13, 14^ This property extends to recognition of synthetic inhibitors, as co-crystal structures reveal that MYST inhibitors engage multiple conserved active contacts, particularly backbone amide hydrogen bonds involved in binding the pyrophosphate moiety of acetyl-CoA (**Fig. 1C-D**).^7, 9, 15^ Biochemical experiments indicate MYST inhibitors target KAT6A/KAT6B over closely-related KAT7 with specificities ranging from 10-to 1000-fold (**Table S1**).^7, 9, 15^ This variable selectivity window is reminiscent of kinase inhibitors, which similarly must differentiate between closely related active sites and often display dose-dependent selectivity.^16^ Direct biochemical and cellular comparisons of sulfonohydrazide and benzisoxazole sulfonamide MYST KAT inhibitors have not been reported. An additional challenge in comparing literature selectivity profiles of MYST inhibitors is that the reported half-maximal inhibitor concentration (IC_50_) measurements arise from biochemical assays whose conditions often differ. In particular, the use of full-length enzymes or reconstituted MYST complexes – in contrast with excised catalytic domains – is uncommon.^12^ The biochemical selectivity of MYST inhibitors within their physiologically-relevant multiprotein complexes remains largely unexplored.

Here we present a comparative analysis of drug-like MYST acetyltransferase inhibitors. First, we use chemoproteomics to validate the ability of the industry-developed KAT6A/B inhibitor PF-9363 to engage MYST acetyltransferases within their native multiprotein complexes. This provides evidence that in addition to KAT6A/B, elevated concentrations of PF-9363 can hierarchically occupy KAT7, KAT8, and KAT5 in the context of their native multiprotein complexes. The ability of PF-9363 to antagonize chemoproteomic capture of MYST complexes, in combination with clustering and structure prediction, is used to identify the transcription factor FOXK2 as a new candidate MYST complex interactor. Our knowledge of PF-9363’s dose-dependent target engagement allows us to validate specific histone acetylation biomarkers for KAT6, KAT7, KAT8, and KAT5, which we apply to study the selectivity of the sulfonohydrazide MYST inhibitors WM-8014, WM-1119 and WM-3835 as well as the mechanisms of acute growth inhibition caused by PF-9363 in the NCI-60 cell line screen. Our studies indicate the highly similar active sites of MYST KAT enzymes render them susceptible to dose-dependent target engagement, similar to how ligands engage the ATP-binding sites of closely related kinases, and highlight general methods to evaluate selective versus multi-enzyme inhibition in proteomic and cellular settings.

## Results

### Chemoproteomics reveals dose-dependent MYST complex engagement by PF-9363

Structural studies indicate MYST inhibitors achieve active site binding by mimicry of the pyrophosphate moiety of acetyl-CoA. However, while the activities of these molecules against purified MYST enzymes in biochemical assays are well-established, their ability to engage native MYST complexes in a proteomic mileau remains uncharacterized. To address this, we applied a chemoproteomic strategy to monitor MYST inhibitor selectivity (**Fig. 2A**).^17, 18^ This method uses a resin-immobilized CoA analogue to afford sensitive capture of KATs and other CoA-binding proteins via their active sites. Pre-incubation of proteomes with active-site competitive small molecules blocks KAT capture in a dose-dependent manner, providing a readout of inhibitor potency and selectivity.^19, 20^ To facilitate MYST inhibitor profiling we optimized our previously reported protocols, leveraging nuclear extracts, capture by an H3K14-CoA bisubstrate, and magnetic bead separation to improve throughput and sensitivity (**Fig. S1**). This enabled enrichment of MYST KATs (KAT5, KAT7, KAT8), associated complex members, and many non-MYST (KAT2A, KAT2B, NAA50, NAA40, NAT10, ESCO1, ESCO2, ATAT1) acetyltransferases (**Fig. 2B-C**). Unfortunately, this method did not sample KAT6A or KAT6B, the most potent targets of PF-9363 in biochemical assays.^9^ Whole proteome profiling of HeLa nuclear extracts did not identify KAT6A/B, suggesting these proteins are present at low copy numbers or poorly extracted from chromatin using traditional fractionation protocols (**Fig. S2, Table S2**). With this potential ‘blind spot’ of our method in mind, we set out to determine the dose-dependent effects of PF-9363. Pre-incubation of nuclear extracts with a low concentration of PF-9363 (0.1 μM) reduced the capture of a single acetyltransferase, KAT7, as assessed by western blot (**Fig. 2D**). Proteomic analysis of capture experiments confirmed KAT7 was the most significantly competed CoA-binding protein under these conditions (**Fig. 2E, Table S3**). This is consistent with prior findings indicating that after KAT6A/B, KAT7 is PF-9363’s second-most sensitive MYST target.^9^

**Figure 2.**
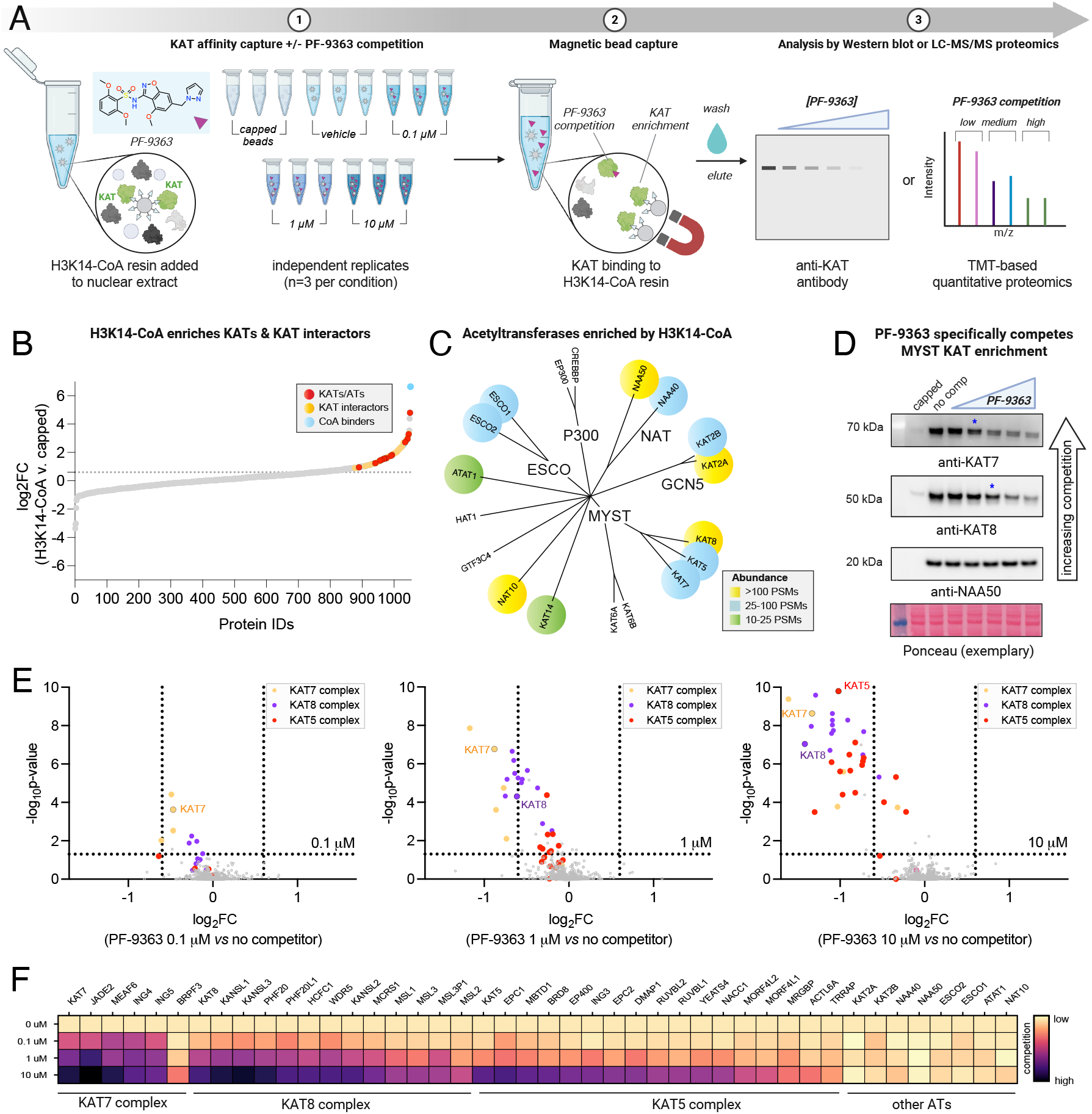
Competitive chemoproteomics reveals dose-dependent MYST occupancy by PF-9363. (A) Schematic of optimized competitive chemoproteomic profiling strategy for KAT enzymes. (B) LC-MS/MS analysis of enriched proteins. Proteins above the dashed line represent targets specifically enriched by H3K14-CoA resin (log_2_FC (H3K14-CoA vs capped)>0.6, p-value<0.05). Red: KATs/ATs; Yellow: KAT interactors; Blue: Other CoA binders; Grey: all other proteins. (C) Phylogenetic tree depicting KATs enriched by H3K14-CoA resin by proteomics. Proteins are color-coded based on peptide-spectrum match (PSM) abundance. (D) PF-9363 dose-dependently competes enrichment of MYST KATs. HeLa nuclear extracts were pre-incubated with escalating concentrations of PF-9363 competitor: 0.01, 0.1, 1, 10, 30 μM (2 h, 4 °C). Ethanolamine-capped beads were used to assess non-specific binding. (E) Proteome-wide competition analysis of PF-9363 competition in HeLa nuclear extracts (*n* = 3 biological replicates). Nuclear extracts were pre-incubated at the specified concentration (2 h, 4 °C) prior to KAT affinity capture. MYST KATs and interactors are color-coded according to complex. (F) Heatmap summary of chemoproteomic competitions of PF-9363 with ATs and associated complex members in HeLa nuclear extracts (*n* = 3 biological replicates). Light colors indicate low engagement, dark colors indicate greater occupancy.

KAT7 operates within a quaternary complex containing MEAF6, an ING protein (ING4/5), and either JADE (1/2/3) or BRPF (1/2/3) subunits.^21^ At low (0.1 μM) concentration, PF-9363 also caused co-competition of JADE2, MEAF6, and ING4/5, indicative of engagement of the JADE complex (**Fig. 2E-F, Table S3**). BRPF3 was also enriched by H3K14-CoA resin, potentially due to capture a BRPF complex. In contrast to the JADE complex, BRPF3 displayed competition only at high concentrations of PF-9363 (10 μM, **Fig. 2F**). We confirmed the differential competition of JADE2 and BRPF3 capture using WM-3835, a putative KAT7-selective inhibitor (**Table S4**). Our chemoproteomic assay cannot distinguish whether this reflects preferential capture of the BRPF complex by the affinity resin, preferential engagement of the the JADE complex by PF-9363, or association of BRPF3 with another MYST complex.

Exploring competition profiles observed at higher PF-9363 concentrations (1 and 10 μM) we find PF-9363 competed capture of all CORUM-annotated members of the KAT8-containing MSL (KAT8, MSL1/2/3, MSL3P1) and NSL complexes (KAT8, KANSL1/2/3, HCFC1, MCRS1, OGT, PHF20, PHF20L1, WDR5)^22^ as well as most members of the KAT5-containing NuA4 complex (MEAF6, EPC1/2, DMAP1, ING3, MBTD1, YEATS4, VPS72, EP400, BRD8, RUVBL1/2, MORF4L1/2, ACTL6A, MRGBP, TRRAP; **Fig. 2F**).^23^ The sensitivity of KAT complexes to PF-9363 follows the general order KAT7 > KAT8 > KAT5. Proteins belonging to multiple KAT complexes are characterized by distinct competition profiles. For example, MEAF6 (KAT5/KAT7) is unique amongst KAT5 complex members in that it also displays potent competition at low concentrations (0.1 μM), presumably reflecting competition of its KAT7 JADE complex-associated fraction. In contrast, TRRAP displays less competition than most NuA4 complex members (e.g. KAT5 itself), reflecting the inability of PF-9363 to compete capture of this protein residing in the KAT2-associated STAGA complex (**Fig. 2F)**.^24, 25^ These studies establish the ability of the benzisoxazole sulfonamide ligand PF-9363 to selectively occupy the catalytic CoA-binding domain of MYST acetyltransferase complexes in complex proteomes.

### Applying chemoproteomic competition to identify new candidate MYST interactors

Encouraged by the ability of chemoproteomics to identify annotated members of MYST complexes, we next sought to assess whether any unannotated proteins displayed similar patterns of PF-9363-mediated competition, potentially indicative of novel interactors. To explore this in a systematic manner we analyzed our chemoproteomic data using an approach similar to our previously reported CATNIP method (**Fig. 3A**).^17^ Briefly, fold-change values generated from capture experiments conducted in the presence of 0, 0.1, 1, and 10 μM PF-9363 were transformed,^26^ plotted in two dimensions, and subjected to k-means clustering.^27^ Five groups of proteins were identified, with two forming a relatively tight cluster and three more disperse (**Table S5**). Inspection of individual proteins within each cluster revealed characteristic dose-response signatures (**Fig. 3B**). The capture of proteins in clusters 2 and 3 were antagonized by PF-9363 in a dose-dependent fashion, with cluster 2 being slightly more sensitive than 3 (**Fig. 3B, top/middle**). Proteins in other clusters, including acetyltransferases such as NAT10, ESCO1, and KAT2B, were relatively insensitive (**Fig. 3B, bottom**). Clusters 2 and 3 encompass 35 proteins in total, 33 of which are known members of KAT7 (BRPF/JADE), KAT8 (NSL/MSL), or KAT5 (NuA4) complexes (**Fig. 3C**). KAT7 JADE complex members were found exclusively in cluster 2, while other MYST complex members were distributed across both. The variant histone H2.AZ (Uniprot: H2AV) was an unexpected cluster member which displays dose-dependent competition by PF-9363 (**Table S3**). Previous studies have found that in addition to histone acetylation, KAT5-containing complexes also mediate ATP-dependent exchange of H2A-H2B for H2A.Z-H2B,^28^ providing a rationale for association with NuA4. The other uanticipated protein found to show a similar pattern of competition to MYST complex members in chemoproteomic experiments was the transcription factor FOXK2 (**Fig. 3C**).

**Figure 3.**
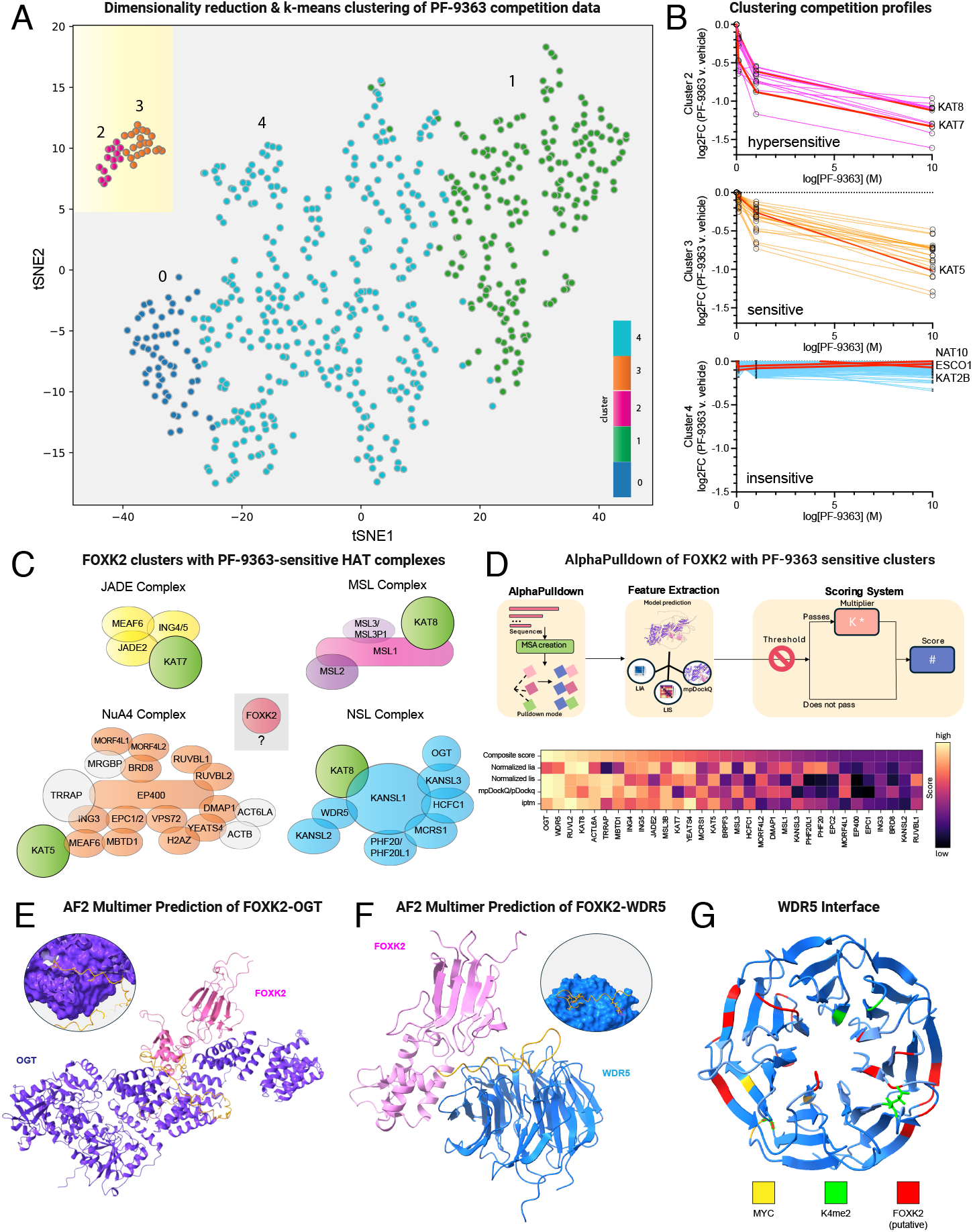
Applying chemoproteomic competition to identify new candidate MYST interactors. (A) t-SNE clustering of proteins based on PF-9363 chemoproteomic competition profile (PF-9363 = 0, 0.1, 1, 10 μM). (B) Dose-response profiles of PF-9363 competition clusters. Colored lines indicate the capture profiles of individual proteins at each concentration of PF-9363 competitor. Red lines represent the capture profiles of labeled KAT or AT enzymes. (C) PF-9363 sensitive proteins identified by this analysis. Green: KAT proteins; Orange: NuA4 complex, Blue: NSL complex; Yellow: JADE complex; Grey: MYST complex proteins not clustered by t-SNE; Red: new candidate MYST interactors. (D) Top: Schematic of interaction screening of FOXK2/PF-9363-sensitive proteins by AlphaPulldown. Bottom: Ranking of predicted FOXK2-interacting proteins by AlphaFold metrics and composite score. Light colors indicate stronger prediction confidence, dark colors indicate weaker prediction confidence. (E) AF3-Multimer predicted structure of FOXK2 (pink) bound to OGT (purple). FOXK2 region predicted to interact with OGT is shown in yellow. (F) AF3-Multimer predicted structure of FOXK2 (pink) bound to WDR5 (blue). FOXK2 region predicted to interact with WDR5 is shown in yellow. (G) Predicted structure of WDR5, highlighting residues known to interact with the transcription factors (MYC, yellow), histones (K4me2, green) as well as predicted FOXK2 interface (red).

FOXK2 is a member of the forkhead box transcription factor family, known for their evolutionary conserved winged helix DNA binding domains and wide-ranging involvement in biological processes.^29, 30^ Circumstantial evidence suggests FOXK2 requires partner proteins to carry out its functions. FOXK2 binds the interleukin-2 (*IL2*) promoter but may enable *IL2* expression by altering the structure of chromatin.^31^ Similarly, genome-wide binding studies indicate association of FOXK2 with forkhead consensus sequences is necessary but not sufficient for target gene expression.^32^ To better understand the ability of FOXK2 to interact with MYST complexes, we performed an in silico interaction screen using AlphaPulldown.^33, 34^ AlphaPulldown is an iterative implementation of AlphaFold2 that uses a ColabFold Search to predict interactions between a single bait protein (e.g. FOXK2) and a large set of potential binding partners, allowing high-throughput discovery of candidate protein interactors (**Fig. 3D**). We applied this to predict binding conformations of FOXK2 in complex with all 34 proteins showing similar chemoproteomic competition profiles (clusters 2 and 3, above) and then scored them based on multiple parameters including ipTM (Interface Predicted TM-Score), normalized LIS (Local Interaction Score) and LIA (Local Interaction Area),^35^ and mpDockQ/pDockQ (**Table S6**).^36^ Additionally, we also introduced a composite score which sums the normalized min-max LIA, LIS, and mpDockQ/pDockQ weighted by a filter based confidence coefficient (Supplementary Information). Ranking these interactors by composite score revealed the top hits to be OGT and WDR5 (**Fig. 3D**). In both cases, interactions were predicted to occur through a region of FOXK2 that was largely disordered (**Fig. 3E-F**). Given the intrinsic uncertainty in predicting interactions in disordered regions, we performed additional analysis of residue-residue contacts using AlphaFold3. This revealed modest but specific contact probabilities with high structural confidence despite weak interaction scores (**Table S6**). In the case of WDR5 the predicted interaction spans surfaces of the protein involved in both histone and transcription factor binding (**Fig. 3G**), with both AlphaFold2 and AlphaFold3 predicting similar interface residues in close in spatial proximity (**Table S6**). STRING analysis revealed OGT has been previously linked to FOXK2 as a member of the polycomb repressive deubiquitinase (PR-DUB) complex.^37, 38^ Direct links between WDR5 and FOXK2 have not been annotated. However, members of the NSL complex (KANSL1/KANSL1L) co-immunoprecipitate with FOXK2,^39^ supporting the ability of this transcription factor to fractionally occupy the MYST complex in pulldown experiments. Future studies will be required to validate the FOXK2-NSL interaction and its biological relevance. These studies demonstrate a strategy for leveraging chemoproteomics and MYST selective inhibitors to identify new candidate interaction partners of multiprotein KAT complexes.

### PF-9363 elicits dose-dependent changes in MYST-regulated histone acetylation

Our chemoproteomic data indicate that high concentrations of PF-9363 are capable of biochemically occupying the active site of several MYST complexes in nuclear extracts. This raises the question: to what extent is this phenomenon biologically relevant (e.g. do high concentrations of PF-9363 hierarchically engage KAT6, KAT7, KAT8, and KAT5 in cells)? To understand how PF-9363 impacts global histone modificatons in a relatively unbiased manner, we profiled its effects across an escalating gradient of concentrations using a bottom-up proteomic method.^40^ In this approach. MCF-7 breast cancer cells are treated with increasing doses of PF-9363 (0.1, 1, 10, 30 μM) or vehicle (DMSO) for 24 hours (**Fig. 4A**). Histones are then extracted from cell pellets and chemically derivatized with propionic anhydride, digested with trypsin, subjected to a second round of propionylation, and analyzed by LC-MS/MS. Measuring the relative intensity of modified versus unmodified forms of individual histone peptides enables simultaneous monitoring of multiple potential biomarkers of MYST inhibition. To minimize false positives, we focused on abundant and reproducible modifications (average stoichiometry of greater than 1% across triplicate samples, standard deviation less than or equal to the average measurement). To account for potential histone modification crosstalk, changes in both acetylated and methylated peptides were evaluated. 38 modified peptides passed this reproducible detection threshold including several established KAT biomarkers^41^ including H3K9Ac (KAT2A/B), H3K18Ac (KAT3A/B), H3K14Ac (KAT7), H3K23Ac (KAT6A/B),

**Figure 4.**
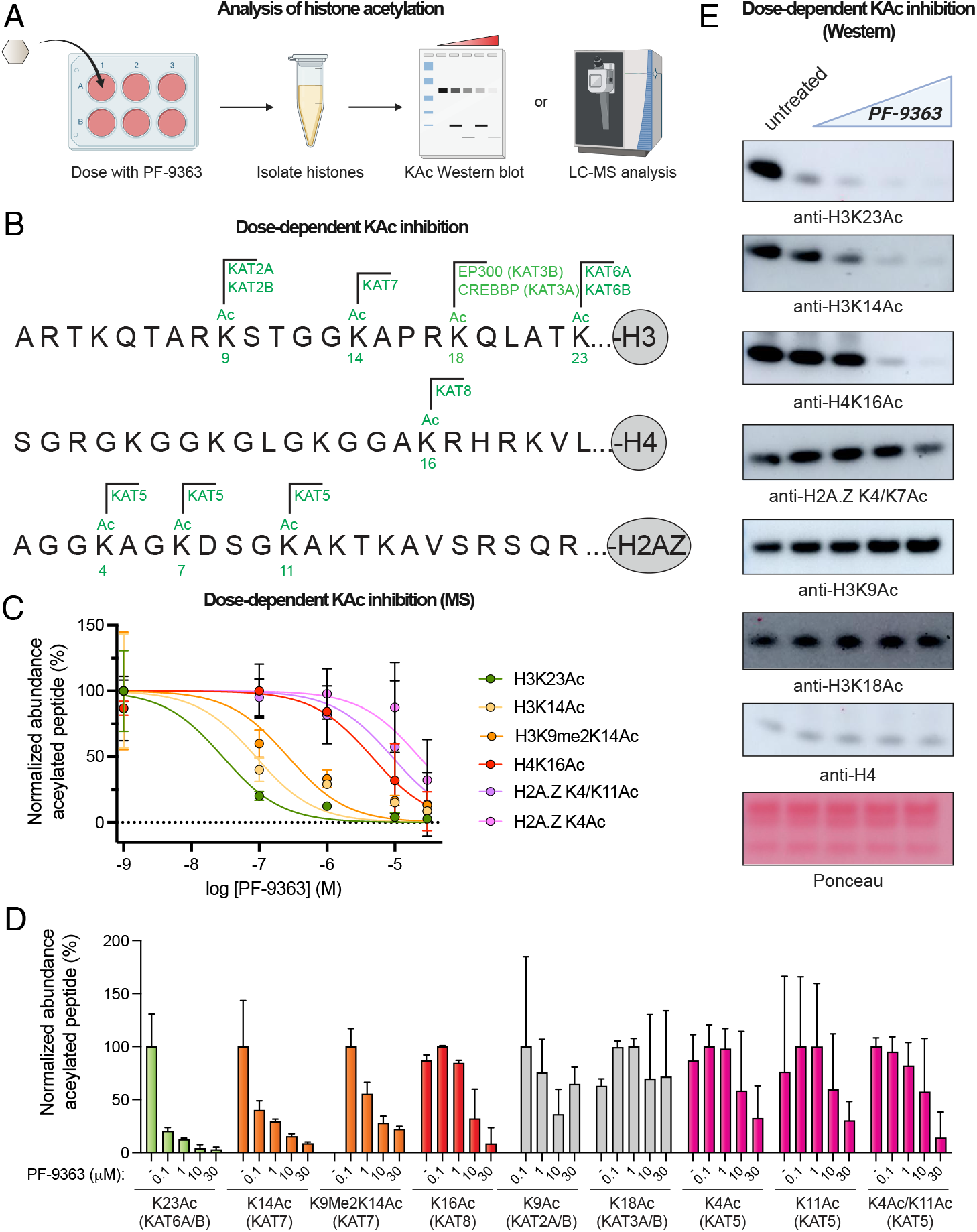
Histone biomarker analysis reveals hierarchical inhibition of MYST acetyltransferases by PF-9363. (A) Schematic of histone biomarker assay used to assess PF-9363 inhibition of KAT-mediated acetylation in MCF-7 cells. (B) Annotated histone acetylation biomarkers of MYST and non-MYST KAT enzymes. CREBBP and EP300 are used interchangeably with their synonymes KAT3A and KAT3B. (C) Dose-dependent inhibition of histone acetylation biomarkers sensitive to PF-9363 as assessed by bottom-up LC-MS/MS proteomics (fold-change ≥ 2, 0 v. 30 μM PF-9363, *n* = 3 biological replicates). (D) LC-MS/MS analysis of histone acetylation biomarkers upon treatment with escalating doses of PF-9363 (n = 3 biological replicates). PF-9363-sensitive markers are colored according to MYST complex; insensitive markers are colored grey. (E) Facilte monitoring of KAT-regulated histone acetylation in response to PF-9363 by western blot (MCF-7 cells, 24 h treatment, 0, 0.1, 1, 10, 30 μM). Data is representative of *n* = 2 biological replicates.

H4K16Ac (KAT8), and H2A.ZK4/K7Ac (KAT5; **Fig. 4B**). Eight peptides displayed a ≥2-fold change in abundance between low-dose (0.1 μM) and high-dose (30 μM) PF-9363 treatment condition, all of which contained acetylated lysine residues (**Fig. 4D, Table S7**). No exclusively methylated peptides were significantly changed by PF-9363 treatment, a phenomenon we confirmed via western blot analysis of H3K27 and H3K79 sites (**Fig. S3**). This is in line with prior genomic analyses of PF-9363, which similarly did not observe changes in the genome-wide distribution of H3K4 or H3K27 methylation.^9^ These studies indicate the acute effects of PF-9363 on abundant histone marks in MCF-7 cells are limited to inhibition of acetylation.

Modified peptides whose abundance was reduced by PF-9363 treatment correspond to six acetylysine positions: H3K23Ac, H3K14Ac, H4K16Ac, H2A.Z4, H2A.Z15, and H2A.Z11 (**Table S7, Fig. 4B**). Dose-response profiling revealed that across these acetylations, H3K23Ac was most sensitive to PF-9363, displaying an ~80% reduction upon 0.1 μM treatment (**Fig. 4C-D**). Genetic perturbations have unambiguously determined the dependence of H3K23Ac on KAT6A/B.^9, 41^ The H3K23Ac biomarker critically complements chemoproteomics by allowing sensitive assessment of KAT6A and KAT6B activity, whose active site occupancy is otherwise difficult to assess. Dose-response profiling confirmed that higher concentrations of PF-9363 inhibit H3K14Ac (IC_50_ ~1 μM) followed by H4K16Ac (IC_50_ ~10 μM), consistent with sequential engagement of KAT7 and KAT8 respectively (**Fig. 4C-D**). PF-9363 was not found to modulate H3K9Ac or H3K18Ac at any concentration, indicating these marks are not relevant biomarkers of MYST activity. This is significant since several studies have used H3K9Ac as a proxy for KAT6A/B inhibition (**Table S7**),^8, 42^ even though its regulation is most commonly attributed to KAT2A/B.^41^ Since LC-MS/MS assays require specialized equipment and technical expertise, we also validated a panel of histone acetylation antibodies that recapitulate the dose-dependent inhibition of MYST targets by western blot (**Fig. 4E**). These studies confirm the dose-dependent pharmacology of PF-9363 and establish a panel of biomarkers for monitoring MYST engagement in living cells.

### Assessing target engagement by orthogonal MYST inhibitor chemotypes

Having established an assay to evaluate the cellular selectivity of MYST inhibitors, we extended it to additional compounds. The sulfonohydrazide-based compounds WM-8014, WM-1119, and WM-3835 are commonly used chemical probes of MYST enzymes (**Fig. 5A**). The first member of this compound to be disclosed was WM-8014, a putative KAT6A/B inhibitor that induces cell cycle arrest and senescence in lymphoma models.^8^ A more bioavailable analogue WM-1119 was subsequently developed, and reported to have increased potency and selectivity for KAT6A/B.^7, 8^ A third sulfonohydrazide, WM-3835, has been used as a chemical probe of KAT7 and phenocopies aspects of its deletion in AML models.^14^ A direct comparison of the dose-dependent effects of these inhibitors across MYST-dependent biomarkers has never been performed.

**Figure 5.**
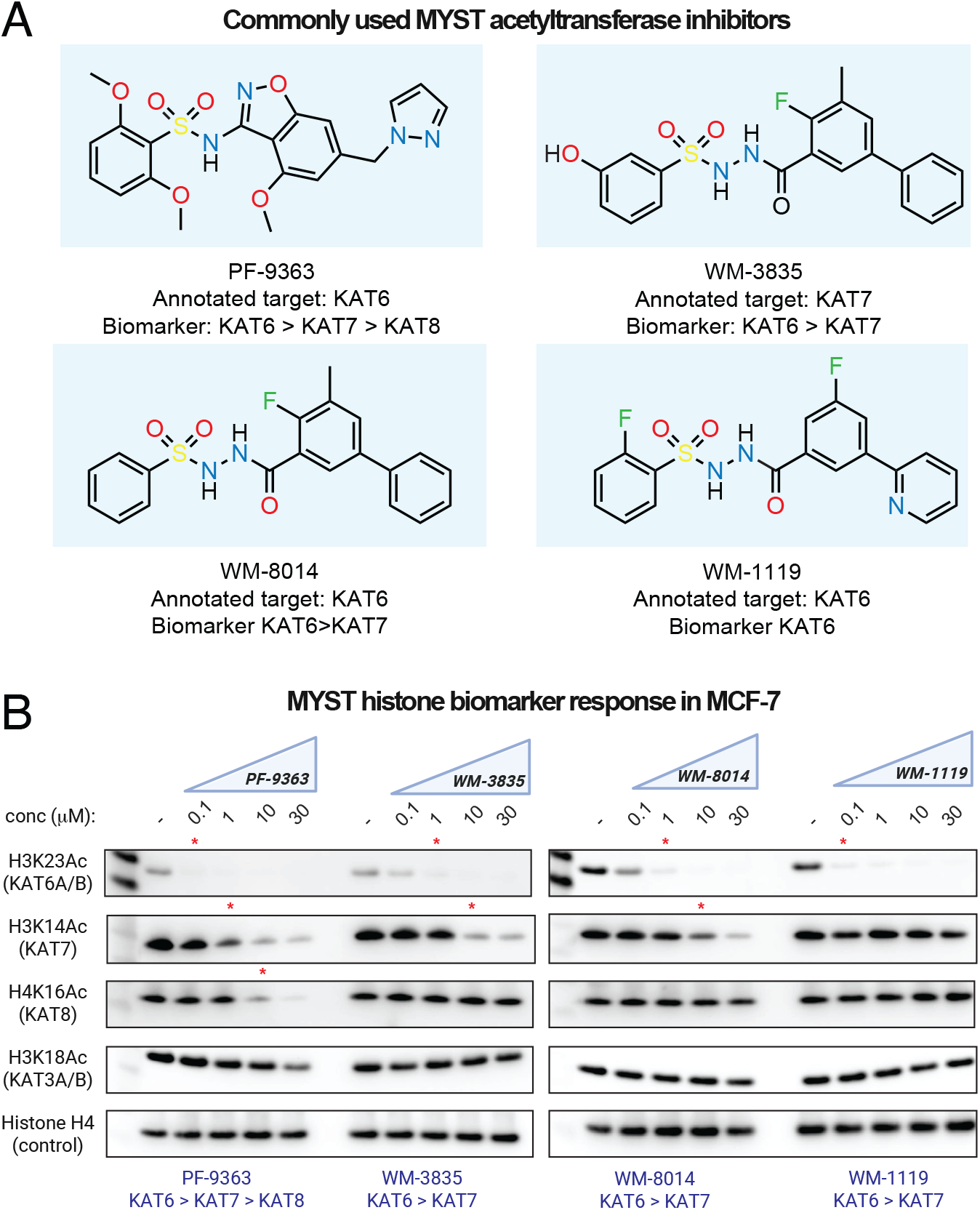
Comparative cellular analysis of orthogonal MYST inhibitor chemotypes. (A) Chemical structures of commonly used MYST KAT inhibitors. Annotated target: KAT enzyme specified as target of each inhibitor in scientific or commercial vendor literature. Biomarker: KAT-regulated histone acetylation biomarker found to be modulated by each compound in this study. (B) Cellular response of histone acetylation biomarkers for KAT6, KAT7, KAT8, and KAT5 to treatment with drug-like MYST acetyltransferases inhibitors. MCF-7 cells were treated with escalating dosages (0.1, 1, 10, 30 μM.) of each compound for 24 h. Red asterisks indicate PF-9363 concentrations where obvious inhibition of a given mark is observed. Data is representative of *n*=2 biological replicates.

To explore this, we treated MCF-7 cells with each compound across a concentration range spanning 0.1 to 30 μM, including PF-9363 as a reference. Target engagement was assessed using the panel of MYST-dependent histone acetylation biomarkers defined above. Analyzing the first generation sulfonohydrazide WM-8014, we observed it was old less potent than PF-9363 but afforded similarly complete inhibition of KAT6A/B – as assessed by inhibition of H3K23Ac – at 1 μM (**Fig. 5B**). At higher doses (10-30 μM) WM-8014 also inhibited KAT7 (H3K14Ac), consistent with prior reports. WM-1119 inhibited H3K23Ac with potency similar to PF-9363, but surprisingly showed little inhibition of KAT7 even at the highest concentrations tested (30 μM; **Fig. 5B**). This observation highlights WM-1119 as a highly selective cellular probe of KAT6A/B activity. While WM-3835 has been applied to study KAT7, its initial publication alludes to its ability to inhibit both KAT6A/B and KAT7.^14, 43^ Consistent with this observation, cellular profiling indicates ~100-fold more potent inhibition of KAT6A/B (H3K23Ac) than KAT7 (H3K14Ac) in the MCF-7 model (**Fig. 5B**). All commercial vendors currently market WM-3835 as a KAT7 inhibitor. Our results highlight a need for caution when interpreting mechanisms arising from WM-3835 treatment and suggest KAT7-dependent phenotypes may be better defined by contrasting the effects of KAT6-selective inhibitors (WM-1119) with dual KAT6/7 antagonists (WM-8014/WM-3835). Overall, these studies demonstrate the how dose-dependent biomarker analysis can inform the application of MYST inhibitors as chemical probes.

### Selective Cytotoxicity of Pan-MYST Inhibition in KAT8-Low Cancer Cells

The effects of MYST inhibitors on cell growth are context-dependent and often require prolonged treatment to manifest. For example, in the initial disclosure of WM-8014, 8 days of compound treatment were necessary for effects on cellular senescence.^8^ Similarly, genetic depletion of KAT6A/B and KAT7 is lethal to ~10-20% of cell lines in the Cancer Dependency Map (DepMap) panel,^44^ but requires 21 days of selection. Extended treatment (21 days) of breast cancer cell lines were used to detect growth inhibition in the initial characterization of PF-9363.^9^ The acute impacts of MYST inhibition on cancer cell growth are less well-characterized.

To explore these effects, we evaluated the sensitivity of the NCI-60 cell line panel to PF-9363 across five doses ranging from 0.1 to 100 μM over 48 hours. As a comparator we used growth inhibition caused by treatment of cells with CPI-1612,^45, 46^ a potent inhibitor of the EP300/CREBBP (KAT3A/B), widely regarded as the master regulator of protein acetylation. Concentrations of PF-9363 expected to primarily effect KAT6 and KAT7 (≤ 1 μM) were generally well-tolerated by the NCI-60 cell line panel, with half-maximal growth inhibition (GI_50_) for most cell lines calculated in the 10-100 μM range (**Fig. 6A, Fig. S4**). In most cases acute inhibition of EP300 and CREBBP by CPI-1612 was much more potent (**Fig. 6A**). An exception to this trend came in BT-549 cells, a triple negative breast cancer model that showed greater growth inhibition at high concentrations (10 and 100 μM) of PF-9363 than CPI-1612 (**Fig. 6B**). Given our prior observation of PF-9363’s unique ability to engage KAT8 and KAT5 at high concentrations, we sought to evaluate if this phenomenon was driving this effect.

**Figure 6.**
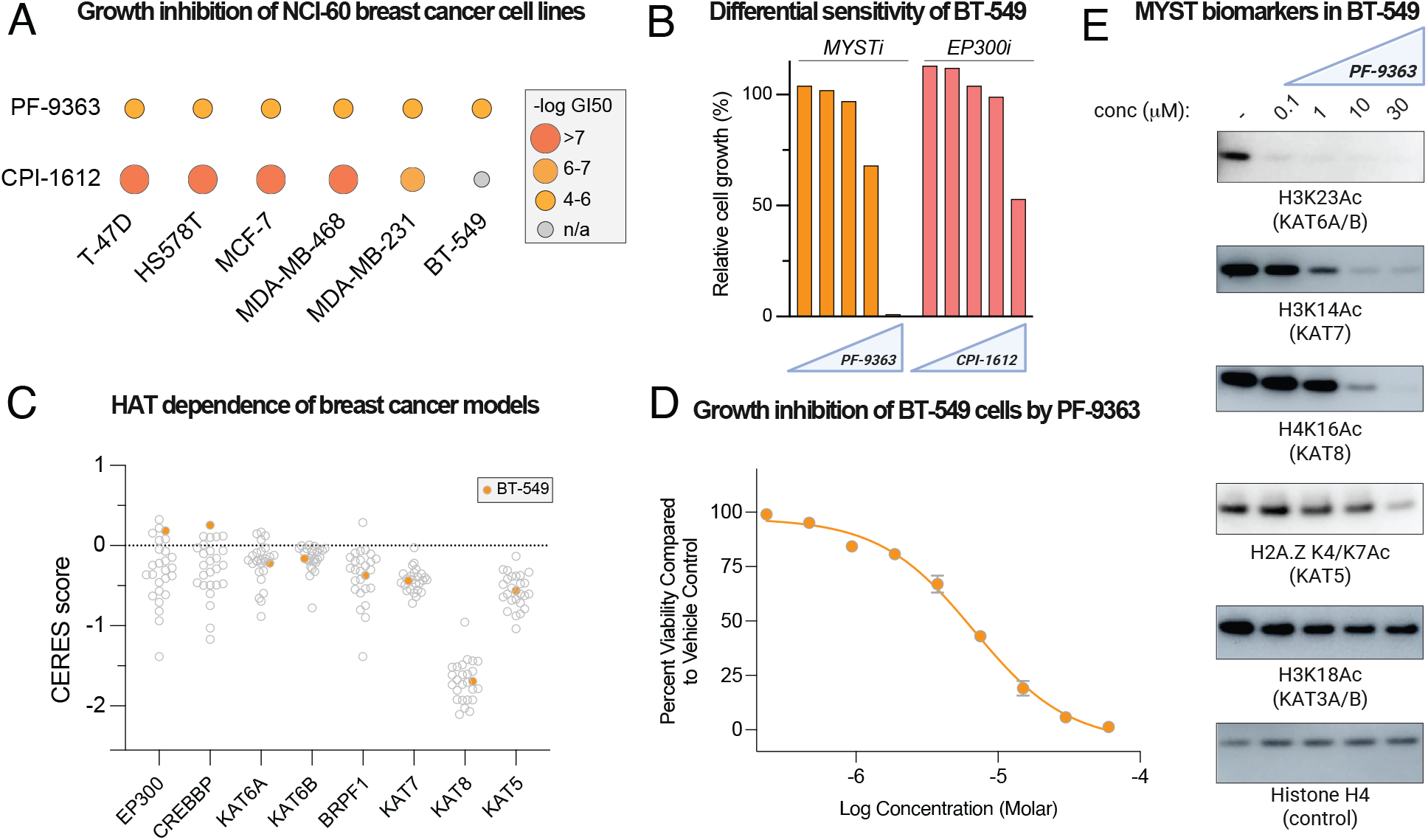
(A) Half-maximal growth inhibition (GI50) values for PF-9363 and CPI-1612 across breast cancer models in the NCI-60 cell line panel (48 h). (B) Relative cell growth of BT-549 cells in the presence of escalating doses of PF-9363 or CPI-1612 (0.01, 0.1, 1, 10, 100 μM, 48 h) as assessed by sulforhodamine B assay in the NCI-60 cell line panel. (C) Dependence of breast cancer cell lines on non-MYST KATs (EP300, CREBBP), MYST KATs (KAT6A, KAT6B, KAT7, KAT8, KAT5), and a MYST KAT complex member (BRPF1) derived from on DepMap CRISPR screening data. BT-549 is highlighted in orange. Lower CERES score indicates a greater dependence of the cell line on the specified gene. (D) Dose-dependent growth inhibition of BT-549 cells by PF-9363 (ATP-Glo assay, 72 h, *n*=4 replicates). (E) Dose-dependent inhibition of of MYST KAT biomarkers by PF-9363 in BT-549 cells (24 h, 0, 0.1, 1, 10, 30 μM). Data is representiative of *n*=2 biological replicates.

Analysis of DepMap-derived CRISPR screening data supports the notion that while breast cancer models vary widely in their requirement for EP300 and CREBBP, BT-549 appears to a fairly independent line. By comparison, targeting of KAT6 and KAT7 exerts modest effects on growth while KAT8 is highly inhibitory, consistent with its designation as an essential gene (**Fig. 6C**).^47, 48^ To validate the NCI-60 results we carried out dose-response cytoxocity assays and determined PF-9363 inhibits BT-549 with a GI_50_ of ~7 μM (**Fig. 6D**). Analysis of histone biomarkers indicated that near complete inhibiton of KAT6A/B (H3K23Ac), KAT7 (H3K14Ac) and KAT8 (H4K16Ac) was achieved near the GI_50_ dose (**Fig. 6E**). KAT3 (H3K18Ac) and KAT5 (H2A.ZK4/K7Ac) were unaffected. Helin and coworkers^49^ have suggested a therapeutic window may exist for targeting KAT8 in KAT8-low tumors, as in these settings KAT8 preferentially associates with the essential NSL complex, whereas in healthy tissues the MSL complex provides a reservoir of KAT8 that can be inhibited without impacting fitness. Evaluation of this strategy may benefit from our observation that WM-8014 and WM-3835 inhibit KAT6 and KAT7 but not KAT8, suggesting their use as controls to define KAT8-independent effects. These studies demonstrate knowledge of MYST inhibitor pharmacology can inform growth inhibition mechanisms and experimental design.

## Discussion

Targeting lysine acetylation via inhibition of KAT activity is an emerging paradigm in oncology,^43^ with several compounds now in clinical evaluation. Critical to identifying new therapeutic contexts for these agents is the proper interpretation of their preclinical effects. Towards that end, here we report a comparative analysis of drug-like MYST acetyltransferase inhibitors. Competitive chemoproteomic profiling of PF-9363 revealed this ligand dose-dependently engages MYST acetyltransferase complexes, but not other CoA-binding sites, in nuclear extracts. This selective engagement allows clustering of chemoproteomic data and can be used to guide de novo identification of known and novel KAT complex members. Integrating chemoproteomic enrichment with AlphaFold screening was used to propose plausible interaction interfaces between FOXK2 and the NSL complex. While validation of this interface was outside the scope of this manuscript, in the future we envision this framework could be useful for the analyzing the composition of KAT complexes in native cells and tissues, as well as be extendable to additional affinity capture matrices.^50^ Unbiased LC-MS analysis of histone modifications recapitulated the hierarchical engagement of MYST acetyltransferase enzymes by PF-9363 as well as known KAT6 (H3K23Ac), KAT7 (H3K14Ac), KAT8 (H4K16Ac), and KAT5 (H2A.ZK4/K11Ac) biomarkers. An exception was H3K9Ac, which in contrast to previous studies^8^ we did not find to be sensitive to MYST inhibition. Given our results and the known regulation of H3K9Ac by KAT2A/B,^51^ we recommend caution when interpreting this modification as a biomarker of MYST KAT activity.

Target engagement data and cellular profiling studies converged on a panel of dose-responsive acetylation biomarkers for each MYST enzyme: H3K23Ac for KAT6A/B, H3K14Ac for KAT7, H4K16Ac for KAT8, and H2A.ZAc for KAT5. Evaluating these marks across multiple MYST inhibitors revealed their surprisingly unique properties. For example, PF-9363 was the only MYST inhibitor that at high concentrations is able to inhibit KAT8-catalyzed H4K16Ac. WM-1119 appears to be a highly specific KAT6A/B inhibitor, leaving other biomarkers untouched even at high concentrations. WM-3835, marketed commercially as a KAT7 inhibitor, potently inhibits KAT6A/B-catalyzed H3K23Ac. As a general rule we recommend monitoring these histone marks when deploying MYST inhibitors in novel systems and/or at escalated doses. This straightforward experiment, which can be carried out using commercial antibodies, provides a powerful measure to ensure that mechanistic effects are properly attributed to a given KAT enzyme’s inhibition.

Evaluation of PF-9363 in the NCI-60 cell line panel indicated that acute inhibition of KAT6 and KAT7 (≤ 1 μM) is well tolerated, whereas growth inhibition at higher concentrations likely reflects KAT8 blockade. As an example, the triple-negative breast cancer line BT-549 - which exhibits minimal dependency on EP300/CREBBP – was sensitive to high-dose PF-9363 (GI_50_ ~7 μM) that coincided with H3K23Ac, H3K14Ac, and H4K16Ac ablation. It has previously been suggested that cell lines with low KAT8 expression may be uniquely vulnerable to KAT8 inhibition.^49^ Our results open the door to testing this hypothesis by exploiting the differential pharmacology of PF-9363 versus WM-8014/WM-3835, the latter which lack activity against KAT8. Another question is whether the hierarchical target engagement phenomenon described here has clinical implications. The phase 1 clinical trial of PF-8144 in breast cancer provided data indicating modulation of the KAT6-regulated H3K23Ac.^10^ Whether KAT7 or KAT8-regulated biomarkers are impacted by any of the MYST inhibitors currently undergoing clinical testing is unknown, but should benefit from the thorough characterization reported here.

During preparation of this manuscript, a study by Perner et al. reported that dual KAT6/KAT7 inhibition using PF-9363 synergizes with Menin inhibitors to overcome primary and acquired drug resistance in MLL-rearranged leukemia.^52^ This timely work illustrates how an understanding of dose-dependent target occupancy can be used to design strategies that rationally leverage MYST inhibitor polypharmacology for therapeutic benefit. By defining the dose-response relationships that govern small molecule engagement of MYST complexes, our studies provide a framework for interpreting mechanism of action, probing MYST-dependent acetylation in diverse settings, and monitoring the selectivity of next-generation KAT inhibitors.

## Acknowledgements

The authors thank Whitney Lieberman (NCI), Diamond Gallimore (NCI), and Krzysztof Krajewski (UNC) for assistance with materials and pilot experiments, Nathan Coussens (DTP) for NCI-60 screening, and Thomas Paul, Oleg Brodsky, and Rhushi Kulkarni (Pfizer) for helpful discussions. Portions of Figure 2A were generated using image templates from BioRender.com under the institutional license belonging to the National Cancer Institute. This work was supported by the Intramural Research Programs of the National Cancer Institute, Center for Cancer Research ZIA BC011488 (J.L.M.) St. Jude Children’s Research Hospital/P22-07377 (B.A.G.) and NIH/P01-CA196539-11 (B.A.G.). This work utilized the computational resources of the NIH HPC Biowulf cluster (http://hpc.nih.gov). In addition, this project has been funded in part with federal funds from the National Cancer Institute, National Institutes of Health, under contract numbers HHSN261200800001E and HHSN261201500003I. The content of this publication does not necessarily reflect the views or policies of the Department of Health and Human Services, nor does mention of trade names, commercial products, or organizations imply endorsement by the U.S. Government.

## Author contributions

Conceptualization: X.C., A.C., J.L.M. Methodology: X.C., A.C., M.P., R.H., K.S., R.K. Software: A.C. Validation: X.C., A.C., M.P., R.H., K.S., R.K. Formal analysis: X.C., A.C., R.H., K.S., R.K., J.L.M. Investigation: X.C., A.C., M.P., R.H., K.S., R.K. Resources: X.C., A.C., M.P., R.H., K.S., R.K. Data Curation: X.C., A.C., R.H., K.S., R.K., J.L.M. Writing - Original Draft: X.C., J.L.M. Writing - Review & Editing: X.C., A.C., M.P., R.H., K.S., R.K., B.G., J.L.M. Visualization: X.C., A.C. Supervision: J.L.M., B.G. Project Administration: J.L.M., B.G. Funding Acquisition: J.L.M.

## Methods

Methods and Supplementary Figures are included as Supplementary Information.

## Data availability

All data, code, and analysis scripts will be submitted to Zenodo upon publication and are currently available in our GitHub repository (https://github.com/castral02/competition_assay). Raw mass spectrometry proteomics files and database search results have been deposited at the ProteomeXchange Consortium (http://proteomecentral.proteomexchange.org) via the PRIDE partner repository with data set identifier “MSV000097704” and password “MYST”. Source Data is provided with this paper as a Source Data file.

