## Supplementary Information for "Hierarchical small molecule inhibition of MYST acetyltransferases"

**Table of Contents for Supporting Information**

| | Page |
|---|---|
| Table of Contents | S2 |
| Supplementary Figures | S2-S5 |
| Materials and Methods | S6 |
| References | S10 |

### Supplementary Figures

A

| Incubation and 1st wash buffer | Aggregation reduction | Adhesion to the tube |
|---|---|---|
| Simple (PBS + protease inhibitor) | aggregate | in wash time |
| Simple + 0.1% NP-40 | aggregate | in 4th wash time |
| Simple + 0.1% NP-40 +remove cell debris | reduced | no |
| Simple + 0.1% NP-40 +200 mM NaCl | reduced | no |
| Simple + 0.5% NP-40 | aggregate | no |
| Simple + 1% NP-40+5% glycerol | aggregate | no |

B

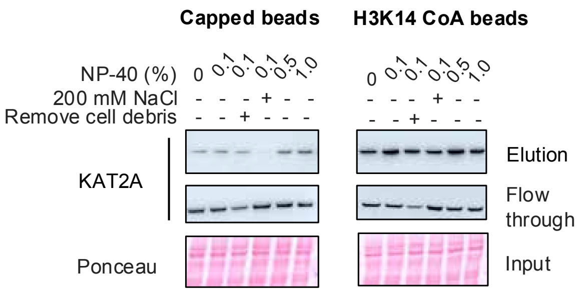


C

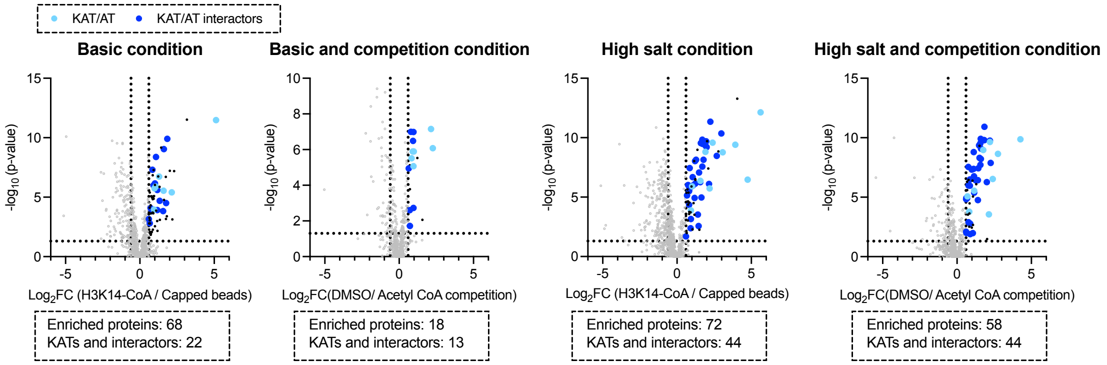


**Figure S1.** Optimization of the chemoproteomic affinity capture assay. (A) Experimental matrix used toe evaluate the effect of incubation buffer and first wash buffer composition on H3K14-CoA magnetic bead pull down assay efficacy. (B) Evaluating the effect of wash buffer on non-specific binding (capped beads) and KAT2A capture (H3K14-CoA beads). For each experiment, HeLa nuclear extracts (200 µL, 1 mg/mL) were centrifuged (20000 g, 30 mins, 4°C) to remove insoluble debris, followed by addition of 10 µL bead slurry and rotation (1 h, 4 °C). Beads were then washed with 500 µL of the indicated buffer and three times with a standard wash buffer (50 mM HEPES pH7.5, 150 mM NaCl, 1 mL). Elution indicates the captured protein eluted from beads, flow through indicates the protein present in the supernatant after incubation with beads. (C) LC-MS/MS proteomic analysis of effect of buffer conditions on KAT capture. Basic condition: PBS, 1x protease inhibitor, 0.1% NP40 (incubation buffer, first wash buffer). High salt condition: PBS, 1x protease inhibitor, 0.1% NP40, 200 mM NaCl (incubation buffer, first wash buffer). Competition conditions refer to the same experiment performed on HeLa nuclear extracts that had been pre-incubated with acetyl-CoA (100 µM, 4 °C, 1 h). Cutoffs of $\log_2$fold-change >0.6 and p-value<0.05 were used to classify proteins as enriched.

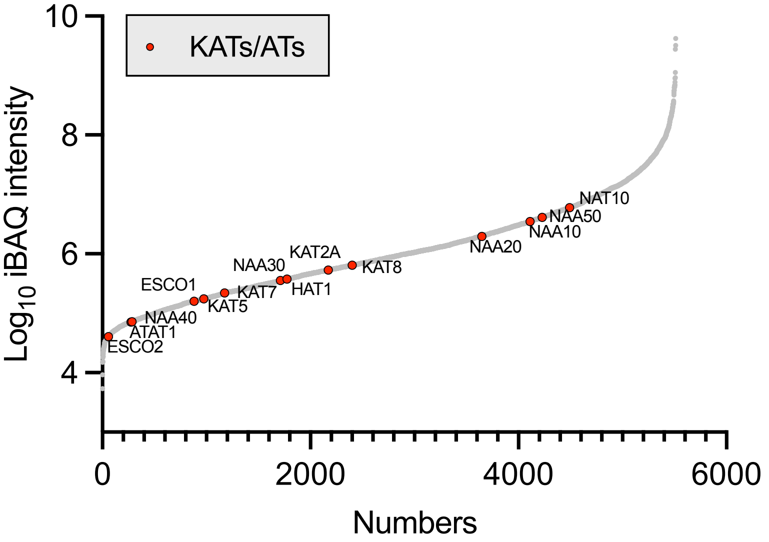


**Figure S2.** Analysis of protein abundance in HeLa nuclear extracts. Samples of nuclear extracts were fractionated, analyzed by LC-MS/MS, and protein abundance assessed by intensity-based quantification (iBAQ) using MaxQuant. KATs and ATs identified from nuclear extracts are colored in red ($n$ = 3 technical replicates). Detailed protein identification data can be found in Table S2.

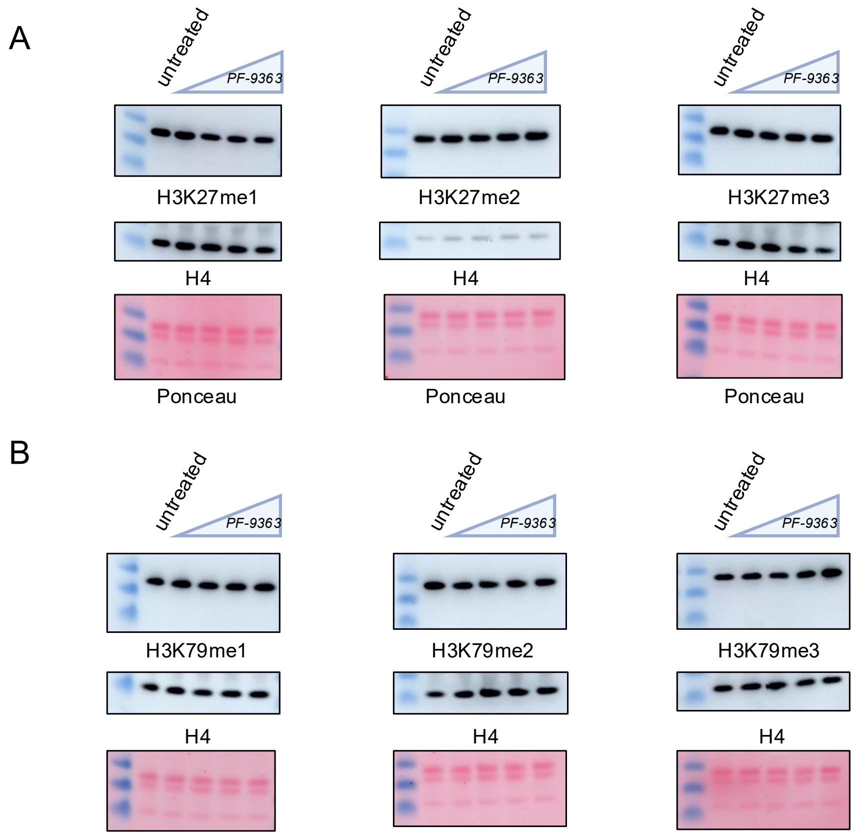


**Figure S3.** Analyzing the effect of PF-9363 on histone methylation. (A) Cellular response of H3K27 methylation to treatment with PF-9363. MCF-7 cells were treated with escalating dosages (0.1, 1, 10, 30 µM.) PF-9363 for 24 h. Data is representative of *n*=2 biological replicates. (B) Cellular response of H3K79 methylation to treatment with PF-9363. MCF-7 cells were treated with escalating dosages (0.1, 1, 10, 30 µM.) PF-9363 for 24 h. Data is representative of *n*=2 biological replicates.

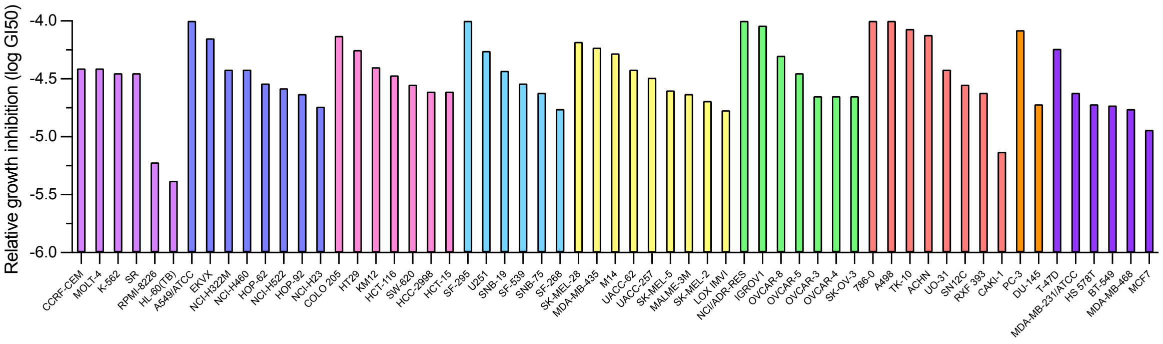


**Figure S4.** Half-maximal growth inhibition ($GI_{50}$).  values for NCI-60 cell line panel treated with PF-9363 (48 h). Lower values correspond to more potent inhibition. Cells lines are colored by lineage as follows: light purple, leukemia; dark blue, non-small cell lung; pink, colon; light blue, CNS; yellow, melanoma; green, ovarian; red, renal; orange, prostate; dark purple, breast.

### Materials and Methods

#### *General materials and methods*

Unless otherwise specified chemicals and solvents were purchased from Sigma, VWR, or Fisher and used without further purification. PF-9363 (HY-132283), WM-1119 (HY-102058), WM-3835 (HY-134901), WM-8014 (HY-102060) and CPI-1612 (HY-136285) were purchased from MedChemExpress. Protein concentrations were determined using the Precision Red Protein Assay (Cytoskeleton, ADV02). Protein samples were prepared for SDS-PAGE by adding LDS buffer (Invitrogen, NP0007) containing 100 mM DTT, followed by denaturation at 95°C for 10 minutes. Samples were loaded onto NuPAGE 4-12% Bis-Tris gels (Invitrogen) and electrophoresed at 160 volts for 1 hour using XCell SureLock Mini-Cells (Invitrogen, EI0002) with MES running buffer (Invitrogen, #NP0002) according to manufacturer's protocols. For detection of MYST proteins, gels were transferred to nitrocellulose membranes using the iBlot dry blotting system (Invitrogen, IB1001) with nitrocellulose transfer stacks (Invitrogen, IB301001) using program 0. For histone acetylation assessment, a modified transfer protocol was employed using the same instrument and transfer stacks but at 20 volts for 8 minutes. Total protein on western blots was visualized with Ponceau staining, followed by two washes with 5% acetic acid in ddH$_2$O. Membranes were blocked with StartingBlock (PBS) Blocking Buffer (Thermo Scientific, 37538) for 30 minutes at room temperature and then incubated with primary antibodies at the indicated dilutions in StartingBlock Blocking Buffer overnight at 4°C. The following primary antibodies were used: anti-KAT7 (Abcam, AB70183, 1:1000), anti-KAT8 (Cell Signaling, 46862S, 1:1000), anti-Naa50 (Proteintech, 16120-1-AP, 1:1000), anti-Lamin A/C (Bethyl Laboratories, A303-431A, 1:2000), anti-H3K23ac (Millipore, 07-355, 1:10000), anti-H3K14ac (Millipore, 07-353, 1:1000), anti-H4K16ac (Millipore, 07-329, 1:1000), acetyl-histone H2A.Z (Lys4/Lys7) (Cell Signaling, 75336, 1:1000), anti-H3K18ac (Millipore, 07-354, 1:1000), anti-H4 (Cell Signaling, 2935S, 1:1000), anti-H3K27me (Cell Signaling, 84932, 1:1000), anti-H3K27me2 (Cell Signaling, 9728, 1:1000), anti-H3K27me3 (Cell Signaling, 9733, 1:1000), anti-H3K79me (Cell Signaling, 12522, 1:1000), anti-H3K79me2 (Cell Signaling, 5427, 1:1000), and anti-H3K79me3 (Cell Signaling, 74073, 1:1000). Following primary antibody incubation, membranes were washed three times with 1× TBST (Cell Signaling, 9997) at room temperature and incubated for 1 hour with either anti-rabbit IgG HRP-linked antibody (Cell Signaling Technology, 7074S) or anti-mouse IgG HRP-linked antibody (Cell Signaling Technology, 7076S), diluted 1:1000 in 1× TBST containing 5% non-fat dry milk. After three additional washes with 1× TBST, membranes were developed using either Lumiglo (Cell Signaling Technology, #7003) or SuperSignal ELISA Femto Substrate (Thermo Scientific, 37074) according to manufacturer's protocols. Images were captured using an Amersham ImageQuant 800 imaging system (Cytiva, 29399482).

### KAT capture and competitive chemoproteomic profiling

#### *Preparation of H3K14-CoA affinity resin and capped beads*

The preparation of magnetic bead H3K14-CoA affinity resin was adapted from a previously reported protocol.[1] Briefly, 15.5 mg (1 eq) of a purified peptide with the linear sequence Ahx-QTARKSTGGK(BrAc)APRKQLATK-NH$_2$ (MW=2260.48 g/mol; UNC High-Throughput Peptide Synthesis Facility) was added to a solution of Coenzyme A sodium salt hydrate (13.9 mg, 2.5 eq, Cayman, 21722) in 750 µL of 100 mM NaHCO$_3$. The reaction mixture was incubated at 37 °C with 300 rpm agitation for 2 hours, then lyophilized. The dried product was reconstituted in 1 mL of 0.1% TFA in water and purified by HPLC using a 100-minute gradient (buffer A: H$_2$O + 0.1% TFA; buffer B: acetonitrile, gradient 25-45% buffer B). The purified Ahx-H3K14-CoA peptide was characterized by LC-MS and lyophilized overnight. For bead functionalization, the Ahx-H3K14-CoA peptide was dissolved in PBS to a concentration of 1 mM. NHS-magnetic beads (550 µL; Pierce, 88827, 10 mg/mL) were pre-washed twice with cold PBS and resuspended in cold PBS to maintain a concentration of 10 mg/mL. The peptide-CoA solution (500 µL, 1 mM) was added to the bead suspension, and the pH was adjusted above 8.0 by adding 12 µL triethylamine. The mixture was rotated at 4°C overnight. The beads were then magnetically separated for 3 minutes, the supernatant was removed, and the beads were washed once with 1 mL $H_2O$. To block unreacted NHS groups, the functionalized beads were incubated with 1 mL of 1 M ethanolamine solution (pH 8.3) using end-over-end mixing for 3 hours at room temperature. The beads were washed three times with wash buffer (100 mM HEPES, pH 7.5, 500 mM NaCl, 1 mL per wash), followed by four washes with IP buffer (50 mM HEPES, pH 7.5, 150 mM NaCl, 0.1% NP-40, 1 mL per wash). The beads were finally resuspended in 500 µL IP buffer to yield a 10 mg/mL suspension and stored at 4°C. Control capped beads were prepared using an identical protocol, except that PBS was substituted for the peptide-CoA solution during the bead coating step.

***Preparation of KAT capture samples for analysis by immunoblotting & LC-MS***

KAT capture and competitive capture using H3K14-CoA magnetic beads, as well as sample preparation for immunoblotting, was performed similarly to previously reported protocols.[1-3] Briefly, HeLa nuclear extracts (IPRACELL, CC-01-20-50) were diluted to 1 mg/mL in high salt buffer (1× PBS [Quality Biological, 114-056-101], 200 mM NaCl, 0.1% NP-40, and protease inhibitor cocktail [Cell Signaling Technology, 5871]). For each assay, 400 µL of diluted nuclear extract was centrifuged (20,000×g, 4°C, 30 minutes) to remove protein precipitates, and the clarified supernatant was transferred to a fresh microcentrifuge tube. Nuclear extracts were pre-incubated with varying concentrations of PF-9363 (0, 0.01, 0.1, 1, 10, or 30 µM) by adding 2 µL of 200× stock solutions in DMSO, followed by rotation at 4°C for 2 hours with end-over-end mixing. After inhibitor pre-treatment, 20 µL of H3K14-CoA magnetic beads was added to each sample, and the mixtures were rotated for an additional 2 hours at 4°C. Beads were collected using a magnetic rack, supernatants discarded, and the resin was subjected to a series of mild washes (1 × 400 µL high salt buffer, followed by 2 × 400 µL standard wash buffer [50 mM HEPES pH 7.5, 150 mM NaCl]). Following the final wash, bound proteins were eluted by resuspending beads in 50 µL of 1× LDS buffer containing 100 mM DTT and heating at 95°C for 10 minutes. After magnetic separation for 3 minutes, the eluted material was transferred to a fresh tube. A second elution was performed using the same conditions, and both eluates were combined. For immunoblot analysis, 20 µL of each elution sample was resolved on NuPAGE 4-12% Bis-Tris gels at 160 volts for 1 hour and transferred to nitrocellulose membranes using the iBlot dry blotting system (program 0). Membranes were probed with antibodies against KAT5, KAT7, Naa50, and Lamin A/C to evaluate the dose-dependent competition effects of PF-9363 on protein binding to the H3K14-CoA affinity resin. Preparation of samples for LC-MS/MS analysis was scaled up 2.5-fold and performed analogously with the following specific modifications: 1000 µL of 1 mg/mL nuclear extract per sample, a modified inhibitor concentration series (0, 0.1, 1, and 10 µM) for PF-9363 and WM-3835, 50 µL of H3K14-CoA beads per sample, proportionally increased wash volumes (1 mL per wash). Instead of LDS buffer elution, beads were resuspended in 50 µL of 50 mM HEPES buffer (pH 7.5) prior to tandem mass tag (TMT) labeling and tryptic digest. Three biological replicates were performed for each condition to ensure statistical robustness. Control experiments using capped beads were processed identically to assess non-specific binding interactions.

### Identification of candidate MYST complex members from chemoproteomic competition data

***Dimensionality reduction and clustering***

Proteomics data containing gene identifiers, log2 fold change values, and associated p-values across three experimental conditions (0.1 v. 0 µM PF-9363, 1 v. 0 µM PF-9363, and 10 v. 0 µM PF-9363) were processed using a custom Python script. Only gene entries with nominal p-values less than 0.05 in at least one condition were retained, ensuring clustering focused exclusively on statistically significant changes. We used log2 fold change values as primary features for clustering analysis. Prior to analysis, entries with missing values were excluded, and features were normalized to zero mean and unit variance. Dimensionality reduction was achieved using t-distributed Stochastic Neighbor Embedding (t-SNE) with perplexity dynamically adjusted based on dataset size (maximum 30) and using two output dimensions. Unsupervised k-means clustering was subsequently applied with the number of clusters set to five. To distinguish between tightly grouped and diffuse clusters, compactness was quantified by calculating the average pairwise Euclidean distance between all points within each cluster. Clusters with lower average pairwise distances were considered more compact, suggesting stronger biological relationships. Results were visualized as two-dimensional scatter plots with points colored according to cluster assignments, revealing distinct gene expression patterns across treatment conditions.

***Structure prediction***

Binding structures between FOXK2 (Uniprot: Q01167) and candidate interactors identified from our chemoproteomic data were predicted using AlphaPulldown[4] v0.30.7,4 with multiple sequence alignments generated via ColabFold Search v1.5.5 on the NIH HPC Biowulf Cluster. Five models were generated per protein-protein pair without using templates or paired MSAs. Local Interaction Scores (LIS) and Local Interaction Area (LIA) were calculated according to previously described formulas.[5] Structural visualizations were generated using ChimeraX v1.8.

***Composite scoring of candidate FOXK2-protein interactions***

To prioritize potential interactions, we developed a composite scoring system integrating multiple AlphaFold metrics (mpDockQ/pDockQ, LIS, and LIA), building upon established approaches for protein-protein interaction analysis.[6] Each interaction was evaluated against minimum acceptable thresholds (1610 for LIA, 0.073 for LIS, and 0.175 for mpDockQ/pDockQ) and assigned a weighted constant (k): 1.0 for passing all three thresholds, 0.75 for two, 0.5 for one, and zero for none. Metrics were then min-max normalized, summed, and multiplied by the weighted constant to generate final composite scores for ranking candidate interactions. All data, code, and analysis scripts can be accessed as described in Data Availability Statement.

***Prediction of interface contacts***

The two highest-ranking interactors by composite score, OGT (Uniprot: O15294) and WDR5 (Uniprot: P61964), were further analyzed to identify interface residue-residue contacts using both AlphaFold2 and AlphaFold3. For AlphaFold2 analysis[7] we calculated Euclidean distances between alpha carbon coordinates of residues using Sci-Py's Distance Matrix Module, filtering for interactions under 8Å. Highly interacting residues were defined as those with distances less than 6Å and PAE scores less than 25.7 For AlphaFold3 analysis[8] we extracted the contact_probs metric from the summary JSON file, with highly interacting areas identified by contact probability values greater than 0.1. Predictions from both methods were compared to identify overlapping interface residues, and structural visualizations were generated using ChimeraX v1.8. All data, code, and analysis scripts can be accessed as described in Data Availability Statement.

**Histone modification analysis**

***Cell culture***

MCF-7 cells were cultured at 37 °C under 5% $CO_2$ in EMEM (Quality Biological, 112-016-101) with 0.01 mg/mL human recombinant insulin (Sigma Aldrich, 91077C), 2 mM L-glutamine (Quality Biological, 118-084-721), and 10% FBS (Avantor Seradigm, 97068-085). BT-549 cells were cultured at 37 °C under 5% CO2 in RPMI-1640 (Quality Biological, 112-024-101) with 10% FBS, 2 mM L-glutamine, and 0.023 U/mL human recombinant insulin.

***Analysis of histone modifications by immunoblotting***

MCF-7 cells ($3.8 \times 10^5$) or BT-549 cells ($3.5 \times 10^5$) were plated per well in 6-well dishes. Experiments were performed in duplicate. After 24 h recovery, media was replaced with 2 mL of inhibitor-containing media to yield final concentrations of 0, 0.1, 1, 10, or 30 µM of the specified inhibitor (PF-9363, WM-8014, WM-1119, WM-3835). Following 24 h inhibitor treatment, cells were washed with ice-cold 1× PBS and scraped in 150 µL of Nuclear Isolation Buffer (NIB) containing 0.1% NP-40 for histone extraction using a modified protocol from the Garcia laboratory. [9] Briefly, cell suspensions were incubated on ice for 5 minutes, and nuclei were pelleted by centrifugation (600 rcf, 4°C, 5 min). Pelleted nuclei were washed twice with 150 µL of detergent-free NIB buffer, followed by centrifugation (600 rcf, 4°C, 5 min) to remove all detergent. Nuclei were then resuspended in 400 µL of 0.4 N $H_2SO_4$ and rotated at 4°C overnight for acid extraction of histones. The following day, samples were centrifuged (11,000 rcf, 4°C, 10 min) and the histone-containing supernatant was transferred to a fresh tube. Trichloroacetic acid (100% TCA) was added to achieve a final concentration of 20% (v/v), tubes were inverted once to mix, and histone precipitation was performed on ice for at least 3 hours or overnight at 4°C. Precipitated histones were collected by centrifugation (11,000 rcf, 4°C, 10 min) and the supernatant carefully removed, leaving a visible film of histones on the tube wall or bottom. Histone pellets were subjected to sequential washes with 1 mL of ice-cold acidified acetone (0.1% 12 N HCl) followed by 1 mL of 100% ice-cold acetone, with centrifugation (11,000 rcf, 4°C, 5 min) after each wash. Samples were air-dried at room temperature, resuspended in 40 µL of $ddH_2O$, and protein concentration determined using the Precision Red Protein Assay (Cytoskeleton, ADV02). For immunoblot analysis, 1-2 µg of purified histones were loaded per lane and processed according to the general immunoblotting protocol described in General Materials and Methods.

***Analysis of histone modifications by LC-MS/MS***

Preparation of histone samples for LC-MS/MS was performed analogously to the procedure for immunoblotting above, with the following modifications. MCF7 cells ($4.2 \times 10^6$) were seeded in 15 cm dishes and treated with PF-9363 or vehicle. Experiments were performed in triplicate. Following treatment and PBS washing, cells were scraped and collected in 15 mL falcon tubes, pelleted (500 rcf, 4°C, 5 min), resuspended in 1 mL cold PBS, transferred to microcentrifuge tubes, and pelleted again before flash-freezing in liquid nitrogen for storage at -80°C. For extraction, cell pellets (~100 µL) were resuspended in NIB buffer at a 1:10 ratio, and cells were lysed with NIB buffer containing 0.2% NP-40 for 8 minutes on ice. Nuclei were pelleted at 1,000 rcf for 8 minutes and washed three times with 500 µL detergent-free NIB buffer. Histones were extracted using 800 µL of 0.2 M $H_2SO_4$, and after centrifugation (3,400 rcf, 4°C, 10 min), supernatants were further clarified at 5,000 rcf before TCA precipitation (33% final concentration). Histone pellets were collected (5,000 rcf, 4°C, 10 min), washed at 7,000 rcf, and resuspended in 100 µL $ddH_2O$. Sample quality was verified by SDS-PAGE with Coomassie Blue staining prior to bottom-up LC-MS analysis.[9] Briefly, histone lysine residues were derivatized using a propionylation reagent (regent-to-sample ratio of 1:2) composed of acetonitrile and propionic anhydride in a 3:1 ratio. The pH of the solution was adjusted to 8.0 using ammonium hydroxide. Propionylation was performed twice, with a 15-minute incubation at room temperature between each round. The samples were then dried on speed vac. The derivatized histones were digested with trypsin at a 1:50 ratio (wt/wt) in 50 mM ammonium bicarbonate buffer at room temperature overnight. Subsequently, the N-termini of the resulting histone peptides were subjected to two rounds of propionylation and dried again using SpeedVac.[10] Peptides were desalted using self-packed C18 stage tips, then dried and reconstituted in 0.1% formic acid. Peptide analysis was conducted using a Vanquish Neo UHPLC coupled to an Orbitrap Exploris 240 mass spectrometer (Thermo Scientific). Samples were maintained at 7 °C in the LC autosampler. Peptide separation was achieved using an Easy-Spray™ PepMap™ Neo nano-column (2 µm, C18, 75 µm × 150 mm) at room temperature. The LC gradient consisted of a linear increase from 2% to 32% solvent B (0.1% formic acid in acetonitrile) in solvent A (0.1% formic acid in water) over 48 minutes, followed by an increase to 98% solvent B over the next 12 minutes, at a flow rate of 300 nL/min. Mass spectrometry was performed using data-independent acquisition (DIA) mode. Each acquisition cycle included a full MS scan followed by 35 DIA MS/MS scans with 24 m/z isolation windows spanning from 295 to 1100 m/z. Full MS scans were acquired in the Orbitrap mass analyzer across 290–1100 m/z at a resolution of 60,000, in positive profile mode, using an auto maximum injection time and an AGC target of 300%. MS/MS data from HCD fragmentation was collected in the Orbitrap. These scans typically used an NCE of 30, an AGC target of 1000%, and a maximum injection time of 60 ms. Histone peptide data were analyzed using EpiProfile 2.0[11]

**Cytotoxicity assays**

NCI-60 cell lines were treated with PF-9363 or EP300/CREBBP inhibitor CPI-1612 at five concentrations (0.01, 0.1, 1, 10, 100 µM) for 48 hours and analyzed by sulforhodamine B assay for growth inhibition as previously described.[12-14] o validate the NCI-60 relative activity trends using an orthogonal method, we assessed PF-9363 activity against BT-549 cells using an ATP-based viability assay. Briefly, BT-549 cells ($3 \times 10^3$ per well) were seeded in white-walled 96-well plates (Corning 3610) and allowed to recover for 24 hours. Cells were then treated with a 10-point dilution series of PF-9363 (60, 30, 15, 7.5, 3.75, 1.88, 0.94, 0.47, 0.23, 0 µM) for 96 hours. Experiments were performed in quadruplicate, and cell viability was measured using CellTiter-Glo® Luminescent Cell Viability Assay (Promega, G7572) according to the manufacturer's instructions. Luminescence signals were recorded on a BioTek Synergy 2 plate reader, and results were normalized to vehicle-treated controls (set to 100%). Half-maximal inhibition values ($IC_{50}$) were determined from nonlinear regression analysis of dose-response curves using GraphPad Prism 9.
